# Inferring absolute counts from proportions by constraining multivariate normal distributions

**DOI:** 10.1101/2025.11.04.686543

**Authors:** Jeffrey Hage, Devin Koestler, Brock Christensen

## Abstract

Biological measurements often result in proportional data, which derive from underlying biological counts. Proportion data are lacking a dimension of information as compared to counts, restricting available analysis methods and separating the data from the biology. We demonstrate a mathematical technique that estimates absolute counts corresponding to proportion data, which we refer to as Mahalanobis Count Inference (MCI). MCI uses information from a population-representative multivariate normal (MVN) distribution of component counts and ultimately outputs an estimated count and a confidence interval per observation proportion vector. We apply MCI to the imputation of white blood cell (WBC) counts, and of total mRNA within single cells. The method performs very well on total mRNA recapitulation (log-space Pearson’s R = 0.81), and well enough on WBC counts to outperform proportions at multiple classification tasks. MCI operates with minimal assumptions, and is applicable to many compositional omics.

## Introduction

Compositional and proportional data are common across scientific disciplines, and particularly so in biological measurements[1]. This includes many sequencing-based applications, such as RNA-seq[2] (in bulk and single cells), DNA sequencing for microbiome analyses[3], as well non-sequencing-based: flow/mass cytometry[4], mass-spectrometry[5], methylation-based cytometry[6], and others. Despite the prevalence of compositional measurements, the underlying biological quantities are almost always unconstrained absolute counts. Compositions also suffer from interdependence, leading to false positives and false negatives in statistical analyses[7][8]. This is because increases and decreases of component counts affect every proportion within an observation. Absolute counts thus better reflect the underlying biology and do not have the interdependency issues of proportions.

When dealing with compositional data, one option researchers have is to implement steps in the wet lab to allow recovery of the absolute scale. For RNA-seq this can be done with known quantities of ‘spike-ins’[9][10], though they are usually used for bias detection and calibration[11]. Conceptually similar absolute quantification methods for flow cytometry are common[12]. For methylation-cytometry, matched complete-blood-counts (CBCs) can be used[13]. Wet-lab methods for microbiome quantification also exist[14]. However, it is not always feasible to implement this step in an experimen-tal plan, and massive amounts of data already e=x2ist where these steps were not taken. The other option is to account for compositionality in data analysis. To allow the data to escape the simplex, researchers can use log ratio transformations[15], such as centered-log-ratio (CLR), additive-log-ratio (ALR), or isometric-log-ratio (ILR)[16]. While these methods can transform data to better fulfill assumptions of many statistical models[1], they do not recapitulate any information about absolute counts, and cannot make compositional data directly comparable to absolute counts[15].

Few non-wet lab methods exist that attempt to recapitulate absolute counts information. Existing methods rely on clever observations about behavior of the specific-omic environments and are restricted to population level inferences[17], or outputs not on the original scale[18]. Methods also exist in mRNA analysis to estimate scaling factors for differential expression analyses, based on assumptions that most transcripts will have similar absolute counts[19], which is analogous to absolute count recapitulation. However none of these apply directly to absolute count inference in the general case of a composition.

Here, we describe an approach to infer the missing absolute count scalar directly from proportions, through constraining the multivariate normal distribution of component absolute counts to an observation’s proportion vector. We refer to the method here as Mahalanobis Count Inference (MCI). MCI can be applied generally to compositional situations where some degree of regularity can be expected within a subset of the underlying populations of components. We demonstrate MCI through predicting white blood cell (WBC) counts, and total mRNA (post-lysis) within single cells.

## Results

### MCI description

To predict absolute counts, we first describe the underlying multivariate normal distribution of a subset of the components of any composition (Figure 1). This single population level distribution is then used in conjunction with a proportion vector of a specific observation to construct a probability density function (PDF) solely in terms of the absolute count scalar. From the observation-specific univariate PDF, we can then derive the most likely absolute count scalar and confidence intervals. This can be done in both log-space and linear-space.

**Figure 1.**
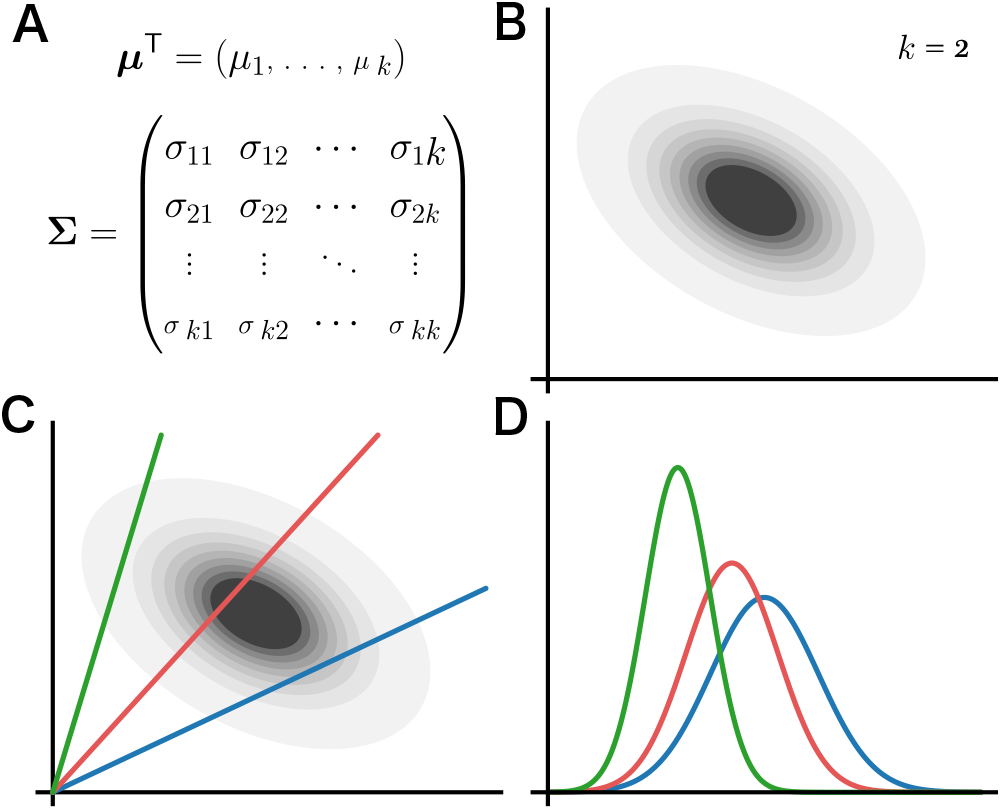
Conceptual overview of Mahalanobis count inference (MCI). **A-B** A multivariate normal (MVN) distribution of component absolute counts. **C** An observation’s proportion vector (colored lines) constrain which counts are possible. **D** A probability density function of possible absolute counts is recovered based on the constraint and the density of the MVN distribution.

For predicting WBC, we applied MCI in four variations, all in log-space. The variations differ in what assumptions are made about the absolute count distributions, and if demographic information is incorporated. **Model 1** assumes that absolute counts of each cell type varies independently. **Model 2** accounts for the covariance of the cell types. **Model 3**, like model 1, assumes that absolute counts of each cell type vary independently. However, it also adjusts the expected counts of each component cell type based on the participant’s age and sex. **Model 4** both accounts for covariance of cell types and adjusts expected counts based on age and sex. For predicting absolute mRNA counts in single cells, Model 2’s structure was used (accounting for covariance in mRNA counts, no demographic adjustment), with an additional feature selection step to select transcripts to include in MCI.

### WBC cohort description

We used data from the 2016 cohort of the Health and Retirement study. Absolute counts of 11 cell types and total white-blood-cell (WBC) counts were derived from CBC and flow-cytometry data. After filtering out all participants with missing and incongruous information, 6,577 observations remained. Of these observations, 3,862 were females (59%), and average age was 68.5 (SD= 10.2) (Table 1).

**Table 1.** WBC cohort characteristics.

| Overall n = 6,577 |  |  |  |
| --- | --- | --- | --- |
| Demographics |  |  |  |
| Age | 68.5 ± 10.2 |  |  |
| Sex |  |  |  |
| Male | 2,715 (41) |  |  |
| Female | 3,862 (59) |  |  |
| Absolute cell counts (×10<sup>3</sup> / μL) |  |  |  |
| WBC | 6.7 ± 3.2 | Eos | 0.21 ± 0.17 |
| Neu | 3.9 ± 1.5 | CD8mem | 0.20 ± 0.18 |
| Mon | 0.56 ± 0.20 | Bnv | 0.097 ± 0.15 |
| CD4nv | 0.42 ± 0.29 | CD8nv | 0.073 ± 0.067 |
| CD4mem | 0.40 ± 0.21 | Bmem | 0.066 ± 2.3 |
| NK | 0.23 ± 0.17 | Bas | 0.056 ± 0.042 |

Continuous: mean ± SD. Categorical: n (%).

We compared how well the frequency distribution of cell counts for each of the 11 cell types and WBC count aligned with normal and log-normal distributions, through comparing maximized log-likelihood. Data aligned better with log-normal distributions for all component counts and WBC count (Supplementary Figure S1, Supplementary Table S1). For this reason, and because the counts of any cell cannot be below 0, all counts were transformed into *log*_2_ space with a small offset (0.003) to account for component observations with some 0 entries (e.g. basophils, eosinophils). All modeling was then conducted in *log*_2_ space. The 6,577 observations were divided into two different sets using age-sex stratified random sampling: 70% (*n* = 4, 595) in the training set and 30% in the testing set (*n* = 1, 982).

### MCI performance on WBC count

Each model was then used to predict WBC count in the training and testing sets (Figure 2). All models outperformed the null distribution by metrics of root mean squared error (RMSE), C-index, and Pearson’s R (all metrics from counts in *log*_2_ space) (Table 2). Interestingly, the four MCI models performed similarly on all metrics with C-indexes near 0.62 for both testing and training sets, indicating that little improvement in prediction accuracy resulted from incorporating covariance and demographic information. Consistency between test and train performance indicates negligible overfitting.

**Table 2.** MCI performance across four models, estimated WBC count compared to true WBC count.

| Model: | One | Two | Three | Four |
| --- | --- | --- | --- | --- |
| **Train** |  |  |  |  |
| Pearson ($\log_2$) | 0.395 | 0.380 | 0.415 | 0.386 |
| RMSE ($\log_2$) | 0.385 | 0.390 | 0.381 | 0.393 |
| C-index | 0.620 | 0.623 | 0.628 | 0.626 |
| **Test** |  |  |  |  |
| Pearson ($\log_2$) | 0.373 | 0.345 | 0.387 | 0.343 |
| RMSE ($\log_2$) | 0.391 | 0.399 | 0.389 | 0.403 |
| C-index | 0.617 | 0.615 | 0.622 | 0.614 |

Pearson is $R$ , not $R^2$ . All metrics on counts in $\log_2$ space. Null C-index is 0.5, null Pearson is 0, and null RMSE is the SD of the count distribution, which is 0.414 for train and 0.418 for test.

**Figure 2.**
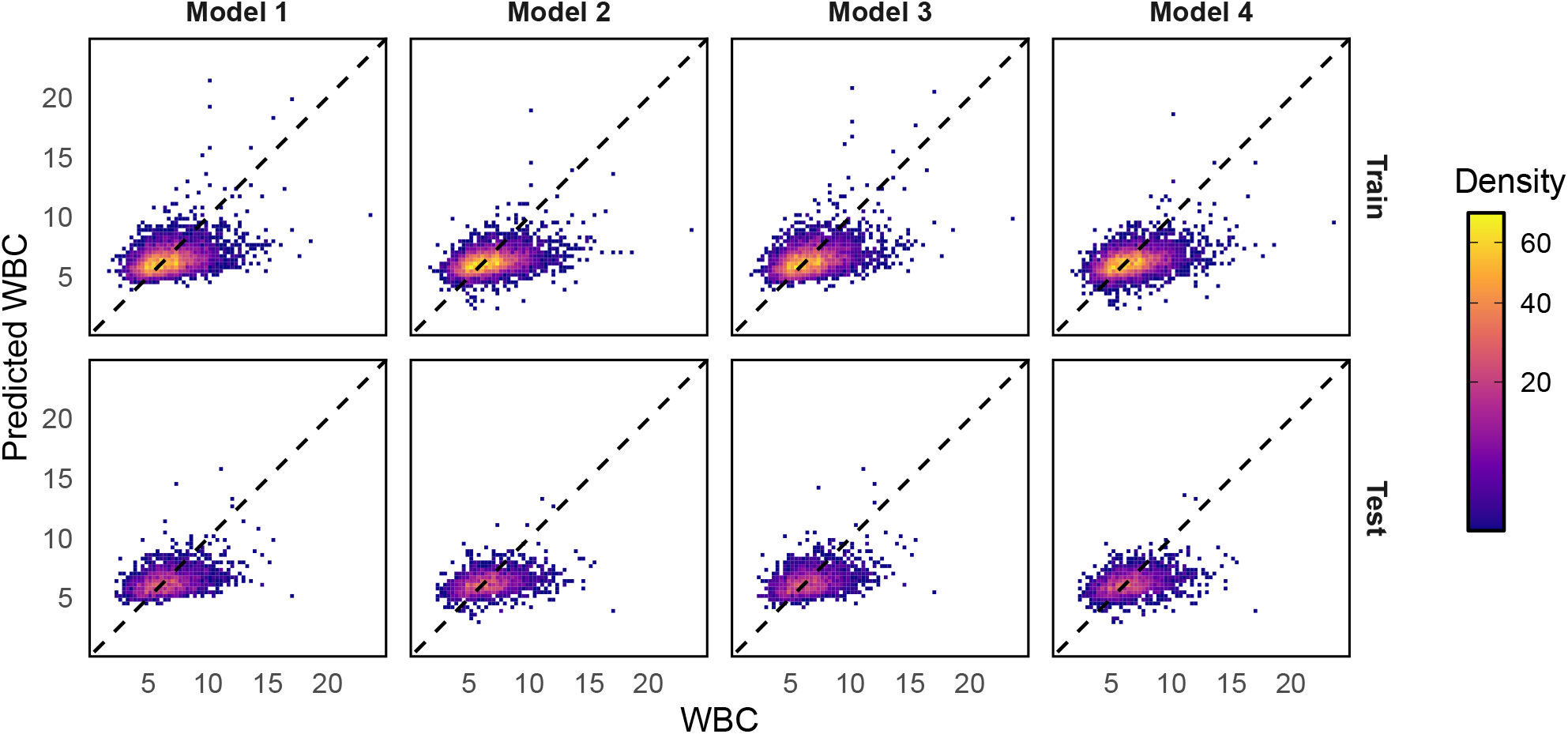
Density plot of WBC count predictions compared to true values across four models in Health and Retirement Study data. Top row is training set, bottom row is testing set. Plots are on a linear scale, in units of 10^3^ cells/*µL*. Dashed line is at unity. Note one observation in each panel was excluded due to being outside view window.

### Confidence interval training

As MCI returns a predicted normal distribution for each proportion vector, confidence intervals can also be derived. The coverage of the 95% confidence interval was assessed for each model, meaning the proportion of true absolute counts that are within the predicted 95% confidence range (Table 3). Model 2 and 4 had much better coverage than models 1 and 3. To account for overdispersion, an empirical multiplier on the predicted log-space standard deviation, *q*, was found for each model to reach 95% coverage on training data. After expansion, all models 95% confidence intervals were similar widths and were accurate on the test set. As shown by low SD-width, confidence intervals in log-space are nearly invariant across observations.

**Table 3.** Confidence interval information for WBC count prediction. All metrics for training set unless specified otherwise. Widths and SD are in *log*_2_ space.

| Metric | Models |  |  |  |
| --- | --- | --- | --- | --- |
|  | One | Two | Three | Four |
| Unadj. Coverage (95%) | 0.82 | 0.94 | 0.79 | 0.93 |
| $q$ expansion for (95%) | 1.49 | 1.04 | 1.60 | 1.12 |
| Mean width (q-adj.) | 1.51 | 1.54 | 1.48 | 1.55 |
| SD width (q-adj.) | 0.02 | 0.02 | 0.02 | 0.02 |
| Test set coverage (q-adj.) | 0.95 | 0.95 | 0.95 | 0.94 |

$q$ multiplies the standard-deviation of outputted prediction in $\log_2$ -space.

### Predicting cell type -penic/-philic

In the setting of compositional cell type measurements such as methylation and flow cytometry (without wetlab spike-ins or matched CBCs) absolute counts are currently underivable. As proportions are compositional, the measured proportions of a cell-type are influenced by the counts of other cell types in the mixture. Thus, we compare the use of our models predicted absolute counts of each cell type at classifying participants into cell-type ‘-penic’ and ‘-philic’ compared to their cell type proportions. The test effectively measures the noise from imperfect absolute predictions against the noise from the compositional environment. The bottom and top 5% percentiles of cell-types true absolute counts were used to assign each of the 6,577 observations to cell-type-penic and -philic. For predicted cell counts, model 2 was used. Among the 11 cell types, the predicted absolute counts significantly (False discovery rate adj. *p <* 0.05) outperformed the true proportions at classifying the cell-type-penic observations in 8 cell types, and the cell-type-philic observations in 8 cell types (Supplementary Figure S2). Across all comparisons, true proportions never significantly outperformed inferred absolute counts.

### Predicting disease status

As diminished or elevated counts of cell types are known to be correlated to a variety of conditions, we tested the ability of estimated cell counts to classify participants in the HRS into disease groups, as compared to true proportions. Model 2 was again used for prediction. Four conditions were selected for measurement due to known cell-type associations and low drop-out: arthritis, diabetes, alcohol-use, and lung-disease. No tested conditions were excluded. The cells used for testing were selected by finding the two highest AUCs of absolute counts into each condition. Among the 8 comparisons, estimated absolute counts had higher AUCs and significantly (DeLong’s test, FDR *p <* 0.05) outperformed true proportions in 5/8 comparisons, and was never significantly outperformed (Supplementary Table S2).

### Inferring total mRNA in single cells

We sought to apply MCI to another compositional setting. Measurements of mRNA abundance via sequencing are compositional. Further, single-cell-omics offer a consistent ‘container’ of the absolute measurement for prediction, making single-cell mRNA prediction a natural fit. We used single-cell data from *n* = 1, 243 naive and primed human embryonic stem cells, grown in vitro. True absolute abundance of mRNA in the cell (post-lysis) was recapitulated using a known quantity of standard External RNA Controls Consortium (ERCC) molecular spike-in for each well. Models in linear-space and log-space were tested and compared, all with a Model-2-like architecture.

### mRNA feature selection

To reduce features included in MCI models from the 964 available features post-filtering, two methods of featureselection were tested. Method 1 greedily minimizes the predicted root-mean-squared error of absolute mRNA count based on the covariance matrix. Method 2, also a greedy selector, empirically maximizes the C-index on the training set via 5-fold cross validation. Note that Method 2 has access to prediction results directly, while Method 1 does not. Each method was tested with each model type, and performance metrics on the training set were gathered with iterative addition of features (Figure 3A-D).

**Figure 3.**
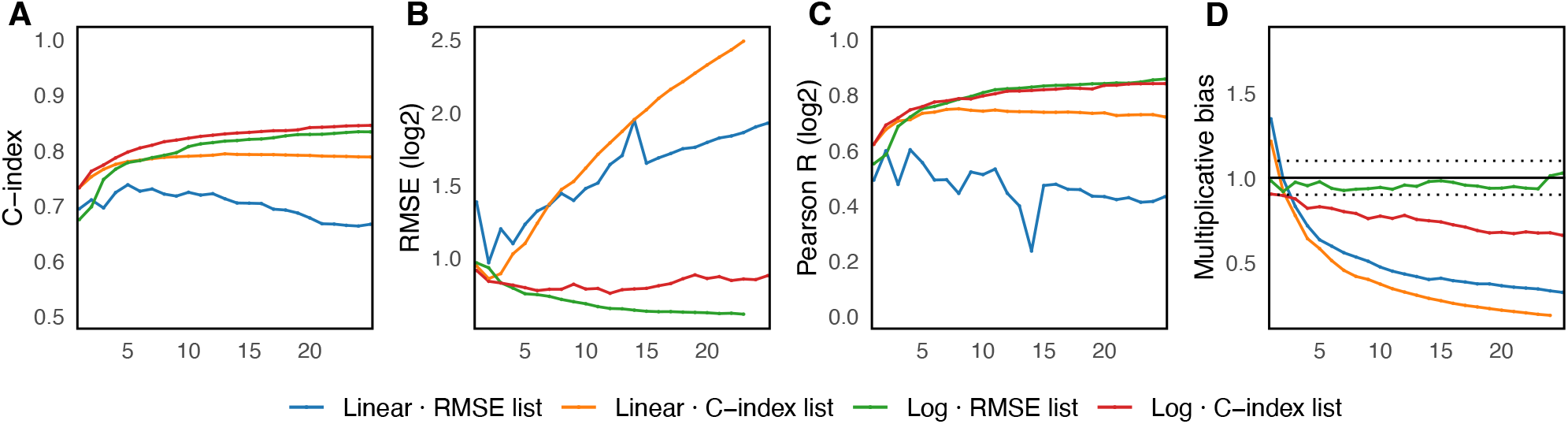
Plots of multiple metrics of mRNA prediction models performance across differing feature amounts (X-axis), on training set. Metrics are: **A** C-index, **B** RMSE (on counts in *log*_2_ space), **C** Pearson’s R (also on counts in *log*2 space), and **D** Multiplicative bias, which is the geometric mean ratio of predicted to actual totals. A value of 1 corresponds to no systemic bias, with dotted lines at *±*0.1. Legend details model and feature list used.

Log-space models outperformed linear-space models on the space-agnostic metric of C-index. Among all model-method pairs, the log-space model with Method 1 feature selection clearly outperformed the others, with nearly the highest C-indexes (next to the greedy C-index log-space model), the smallest RMSE, the highest Pearson R (again alongside the other log-space model), and the most bias free predictions on multiplicative scale. Method 1 also is non-empirical, and purely derived from the MVN distribution.

Due to its superior performance, the log-space with Method 1 was chosen to predict total mRNA on the test set. Based on the analyzed metrics in Figure 3, 15 features were used. Selected features, in order, were: *COX7C, SUMO2, IPO5, HSPA4, SNRPE, HSBP1, RPL31P2, NRDC, LRPPRC, RPL15, HSP90AB1, SRP9, TMSB4X, KDM5B*, and *EIF3M* (Supplementary Table S3). This list encompasses genes from multiple core-biological pathways, as the RMSE method effectively prioritizes stability and independence of features.

### mRNA model performance

The log-space model with the RMSE-minimizing feature selection (Method 1) and 15 features was used to predict total mRNA count on the test and train set (Figure 4). On the testing set, the model-method pair retained a high C-index of 0.802 and a Pearson R (in *log*_2_ space) of 0.810 (Table 4). RMSE in *log*_2_ space was 0.639, and multiplicative bias was near 1.00 (0.982).

**Table 4.** MCI model performance on total mRNA in single cells (top 15 RMSE features in log-space).

| Metric | Train | Test |
| --- | --- | --- |
| C-index | 0.819 | 0.802 |
| RMSE ($\log_2$) | 0.633 | 0.639 |
| Pearson $R$ ($\log_2$) | 0.832 | 0.808 |
| Multiplicative bias | 0.982 | 0.982 |

Pearson is $R$ , not $R^2$ . All metrics in $\log_2$ space.

**Figure 4.**
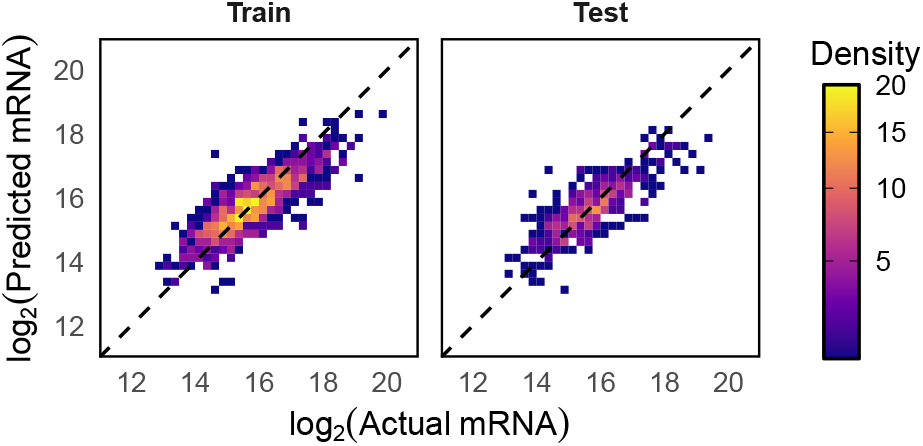
Actual vs. predicted mRNA in single cells in *log*_2_ space, using the top 15 features from RMSE selection method (Method 1) with log-space MCI model. Dashed line is at unity.

## Discussion

### Overview

Here, we developed and applied a mathematical method based on proportion vector constraints along the multivariate normal (MVN) distributions of component counts, allowing recapitulation of absolute scale information. The model, named Mahalanobis Count Inference (MCI), can be applied with both linear and logarithmic scales of component counts and across many biological and non-biological situations. To our knowledge, no similar method has previously been developed or applied to the general case of inference of absolute counts from proportions without wet-lab intervention.

We applied MCI to infer absolute white blood cell (WBC) counts to a degree sufficient to outperform proportions at predicting disease states and low cell-specific counts. We also applied MCI to predicting total mRNA (post-lysis) of single cells, where it performed strongly as quantified by the (logspace) metrics of Pearson’s R, RMSE, and C-index.

### MCI Assumptions and appropriate use-cases

MCI models in linear and log-space both assume component counts follow a MVN distribution in the respective space. For most use-cases the log-space model will be more appropriate as both component counts and the resulting prediction cannot be negative, whereas the linear model leaves non-zero probability density at negative component counts.

A key ingredient is also ensuring that the count being estimated is from a consistent kind and scale of sample. For instance, single cells naturally create a sensible ‘container’ in which to estimate mRNA count. Bulk samples not taken with some consistent size specifications would violate this requirement.

For transportability, it is important that the training situation is directly applicable to the implementation situation of an MCI model. For instance, application of the single-cell mRNA model here may under perform or have systemic bias if applied to other cell types, since they were not represented in training data, and thus their component counts of selected features may not follow the same counts. In the case of WBC, as age is known to be a contributor to the expected WBC-composition, the training data should match the application set by that, and possibly other, demographics. Known observation attributes that may change the expected MVN distribution of component counts can be used to adjust it per-each observation, as demonstrated with age[20] and sex[21] on component white blood cell counts.

For an MCI model to be trained, there has to be absolute counts data of proportion components available, which may not be the case for all compositional situations. Given that absolute counts data is available, MCI is only useful when such data is unavailable in the application environment. This makes MCI apt for uses such as RNA quantification, where spike ins can be used for absolute measurements[10], or for blood profiles, where absolute count information is unavailable in some flow cytometry pipelines, as well as methylation-cytometry based applications without matching CBC or blood smear.

### Feature selection in high-dimensional environments

In the WBC-environment, available features were naturally limited to the cell-types present (and discerned in this context) in blood. However, many different mRNAs are expressed within cells, and thus feature selection was necessary. The log-space model with the predicted RMSE minimization based feature selection performed convincingly against its empirical counterparts, despite not having access to any resulting predictions. This is in line with maximizing Fisher information of the MVN distribution. If MCI is applied to other omics, this method of feature selection has theoretical backing corresponding to the models mechanism, and does not require the predictions themselves to calibrate.

## Methods

### Derivation of MCI in linear-space

Given a situation where a proportional observation is available, we seek to estimate the underlying total count giving rise to the observed proportions. We do so by incorporating information about the underlying MVN distribution of the absolute counts of some subset of the components of the composition in question. Our goal is to construct a univariate-Gaussian probability density function of the absolute count scalar *c*, for an observed proportions vector **P***∈* [0, 1]^*k*^ of dimension *k*. **P**_*i*_ is the proportion of feature *i*. Note that as only a subset of the components in the composition must be included for the inference, the sum of the components of the vector is less than or equal to one, not necessarily exactly one:

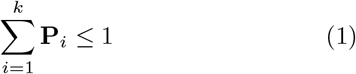

We will first describe this inference in linear-counts-space, then in log-counts-space.

We begin with a MVN distribution (**X**) of component absolute counts across *k* dimensions, shown in Equation (2). The distribution is described by a mean vector ***µ*** *∈* (0, *∞*)^*k*^, and a covariance matrix **Σ** *∈* ℝ^*k×k*^, which is positive-definite and symmetric **Σ** *≻* 0. Note that the population level parameters **Σ** and ***µ*** are taken to be known rather than estimated.

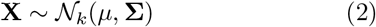

The probability density of a *k* dimensional point **x** can be found easily by finding the squared Mahalanobis distance from the mean vector:

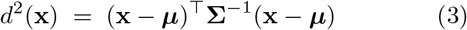

The distance can then be input into the probability density function:

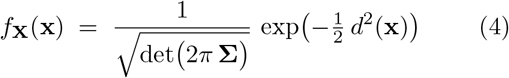

Having a known observation of proportion vector **P** allows us to constrain the possible points in this MVN distribution to those that lie along **x** = *c***P**, where *c* is the positive absolute-counts scalar of interest *c ∈* ℝ_+_. The possible Mahalanobis distances with a known **P** are shown in Equation (5). These distances are only a function of scalar *c*, as all other variables are taken as known and constant.

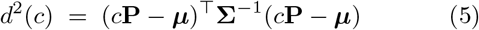

Using bilinearity, the symmetry of **Σ**^*−*1^, and then completing the square in *c*, Equation (5) can be rewritten as:

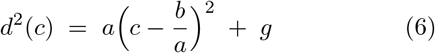

where *a* = **P**^*⊤*^**Σ**^*−*1^**P** *>* 0, *b* = **P**^*⊤*^**Σ**^*−*1^***µ***, and 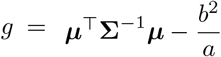

This constrained squared Mahalanobis distance function can then be used to construct a probability density function analogous to Equation (4), however it must be renormalized. Including the unknown normalization constant *K*, which incorporates the constant multiplicative term 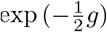, produces:

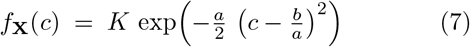

To find the proper normalization constant *K*, the integral in Equation (8) can be solved for *K* through u-substitution into the form of a Gaussian-integral.

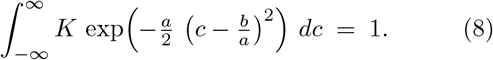

The solution allows construction of the final probability density function of *c* given **P**:

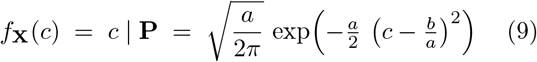

This result is a normal distribution with mean 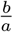 and variance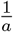 Substituting back in *a* and *b* gives:

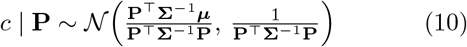

The estimated absolute count scalar *c*č is then simply the expected-value of the normal distribution, and the 95% confidence interval is derived from the variance:

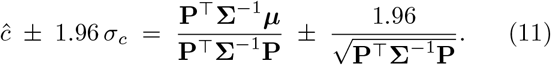

Note that the population parameters **Σ** and ***µ*** are treated as known, though actual uncertainty in parameter estimates may cause overdispersion.

### MCI in log-space

As counts cannot be negative, modeling the underlying component count distributions and absolute count scalar *c* through log-normal distributions may provide better representations. Similar concepts from the previous derivation can be applied with some adjustments. Let **Σ**_***ℓ***_ *∈* ℝ^*k×k*^ represent the covariance matrix of the MVN distribution **X**_*l*_ of the *log*_2_ transformed *k* component counts, and ***µ***_***ℓ***_ *∈* ℝ^*k*^ represent the mean vector. In practice, small offsets would need to be added in this transformation if any 0 counts are present.

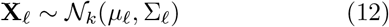

Analogous to Equation (5), the scalar *c* times proportion vector **P** can be inputted into the corresponding squared Mahalanobis distance function, though they must be transformed to log-space. A *k* dimensional vector *ϵ* of a small positive offset can be added to *c***P** to ensure no zero proportions exist:

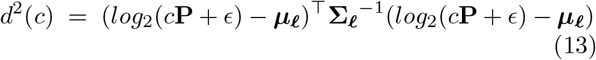

When *ϵ* is the zero vector, then *log*_2_(*c***P**) separates to *log*_2_(*c*)**1** + *log*_2_(**P**), where **1** is the vector of ones. With the replacement *t*:= *log*_2_(*c*):

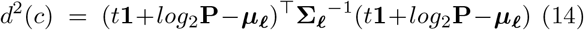

By employing the same procedure in the previous section: rearrangement into the quadratic form of *t*, inserting into the MVN probability-density function, and renormalization, the resulting PDF of *t* given **P** with *ϵ* = 0 is normal:

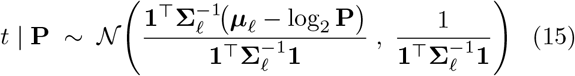

As a consequence of log-space, when *ϵ* = **0** the variance does not depend of the proportion vector, and thus the same confidence interval in log-space is returned for all observations. The estimated *log*_2_ absolute count 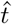 is the expected-value of the normal distribution *t* **P**, and the 95% confidence intervals derived from the standard deviation.

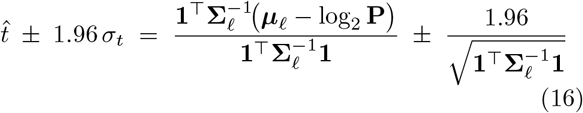

The estimation of the absolute count is then 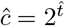and confidence intervals are similarly transformed:

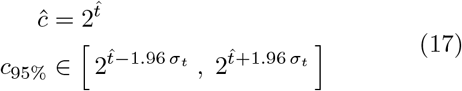

In the case where *ϵ* ≠ **0**, which is the case for predictions implemented here, the cross section of the MVN is not a straight line, which results in a non-normal *t*| **P**. Rather than an analytic solution, 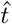 can be estimated numerically through maximizing log-likelihood of *t* for corresponding count points *log*_2_(2^*t*^**P** + *ϵ*) across the MVN distribution. Confidence intervals can be estimated via curvature of log-likelihood *ℓ*(*t*) at 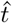.

### Predicting accuracy limits

The accuracy of MCI is limited by the underlying MVN distribution of component absolute counts. In cases where a wide array of component combinations produce similar proportions, it is difficult to discern which absolute scalar *c* corresponds to an observation. This is quantified neatly in the log-space model (assuming *ϵ* = **0**), as the root-mean-squared-error (RMSE) of the log-space prediction of the model 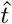 is equivalent to the standard deviation of the prediction (assuming no systemic bias), which is independent of observation’s proportion vector **P**:

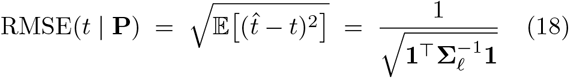

This allows for estimation of MCI performance at the point of log-space MVN construction. For the linear-space model, the same concept applies, though the RMSE in linear-space is dependent on the observation vector **P**. As **P** does not necessarily have components summing to one, the distribution of **P** vectors is not fully determined by the MVN distribution.

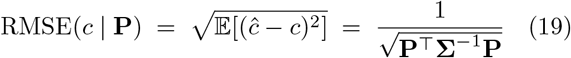

### WBC data source and preprocessing

Absolute counts data of 11 white-blood cell types was sourced from the 2016 cohort of the Health and Retirement Study (HRS). As described previously[22], peripheral blood samples were collected from participants in their home. Among other measurements on the collected samples, a Complete-Blood-Count (CBC) with differential was performed, alongside many-antibody flow-cytometry on cryopreserved mononuclear cells.

For this study, the participants with matched CBC, flow-cytometry, and demographic data were considered. Participants with incomplete data were filtered out. Absolute total white blood cell (WBC) counts were taken from CBC data. 15 rows where CBC proportions didn’t sum to near one (0.97-1.03) were filtered out. Normalized percentages of Neutrophils, Basophils, Eosinophils, and Monocytes were multiplied by WBC counts to retrieve absolute counts. Absolute counts (originally calculated in the HRS through multiplication of flow-cytometry proportions with appropriate CBC counts) of B-naive, CD4-naive, CD8-naive, and Natural-Killers were used directly from reported data. B-memory, CD4-memory, and CD8 memory were derived by summing underlying gate counts, as described in supplementary information (Supplementary Table S4).

One case where the sum of component counts was higher than WBC count was filtered out. 12 cases where WBC count was more than 30% higher than the sum of component counts were filtered. Proportions were taken as count of cell type divided by total WBC count. All considered proportions need not sum to one for an observation, as other populations (such as dendritic and T-regulators) are not represented. All counts and proportions were rounded to three significant figures.

After all filtering, 6,577 observations remained. Observations were split 70-30 for training and testing sets, accounting for balanced splitting of age and sex groups. Observations were stratified into 8 quantiles of age, and between sexes. Random sampling into the training and testing set took place within each sex stratified quantile. The training set had 4,595 observations and the testing had 1,982.

### WBC model implementation

For predicting WBC, the log-space MCI model was used, as counts cannot be negative, and log-normal distributions fit the components counts better than linear-normal (by log-likelihood). However, as the *ϵ* offset could not be **0**, as some components of the proportion vector **P** may be 0 for some samples, the estimation of WBC for each observation was not derived analytically. Instead, the estimation of WBC in *log*_2_ space 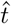 was found through numerical maximization of log-likelihood within the MVN distribution along the constraint defined by the proportion vector *P*:

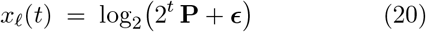

The log likelihood to maximize is then:

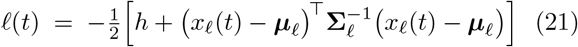

where *h* = *k* log(2*π*) + log |**Σ**|_*ℓ*_ is a constant. As the analytic solution of the 95% confidence interval of 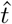, which is independent of **P**, is compromised due to *ϵ* ≠ **0**, the confidence interval is also estimated numerically. This is done by assuming a normal curve, and finding the second-derivative of the log-likelihood function which approximates the standard deviation of *t*:

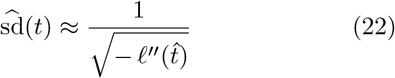

which can then be used to calculate 95% confidence intervals.

Four variations of MCI models were tested for WBC prediction. Model 1 assumes that all cell types vary independently, as implemented through setting all nondiagonal entries in the MVN distribution to 0. Model 2 is the generic model, where the covariance matrix is unstructured. Model 3 is similar to model 1, as it also assumes component count independent by setting all non-diagonal entries to 0. However, model 3 also adjusts the mean vector in log-space *µ*_*ℓ*_ for age and sex. Each component of the *k* dimensional *µ*_*ℓ*_ was modeled with natural cubic splines with 4 degrees of freedom, interacting with sex. This effectively makes the mean vector *µ*_*ℓ*_ a smooth function of age for each sex. Model 4 models the mean vector *µ*_*ℓ*_ as age-sex dependent in the same way as model 3, and also allows an unstructured covariance matrix. Component variances and covariances are not modeled as a function of age-sex, which allows constant multiplicative variance depending on the mean. For Model 3 and 4, variance and covariance (for model four) was estimated through residuals from splines, rather than marginally.

Post initial model fitting on training data, confidence intervals were expanded empirically to encompass 95% of data through finding necessary expansion parameter *q* applied for predicted standard deviations in log-space.

### WBC downstream analyses

Two analyses were done on predicted WBC counts (from model 2) to test the ability of cell-count component predictions to compete with true-proportions at inference of various observation qualities. Both analyses used all 6,577 observations available (post-filtering) from the HRS data.

Observations in the HRS data were classified into cell-type-penic and cell-type-philic for each cell type, based on the bottom 5 percentile and top 5 percentile of each true absolute cell count. This was done as standard penic and and -philic reference ranges are not commonly accepted for all 11 cell types in our data. AUC for each cell type was calculated for both the corresponding true cell proportion, and the estimated absolute count for that cell type. Significance was calculated through a paired, non-parametric DeLong’s test, and comparisons were false discovery rate adjusted (FDR) using Benjamini-Hochberg within -penic and -philic groups separately.

Four disease states were also selected from the HRS data for a similar prediction comparison. Disease states were selected based on known cell-type associations, and low non-response numbers within the HRS 2016 cohort. The four selected states were: Arthritis, diabetes, alcohol use (ever used), and lung disease. No additional disease states were tested and excluded. For each disease state, two cell types were selected for comparison, based on the highest AUCs using true absolute counts, which indicates some classification ability from those cell types. Similarly to the previous analysis, AUC of estimated count was tested against true proportion for each cell-type-disease state pair. Significance was measured by paired, non-parametric DeLong’s test, FDR adjusted across all 8 comparisons.

### Single-cell mRNA data source and preprocessing

Single-cell transcriptomic data was sourced from E-MTAB-6819 on ArrayExpress. As described on the source study protocol [23], naive and primed human embryonic stem cells (WA09-NK2) were proliferated invitro and sorted into individual wells. Importantly, an External RNA Controls Consortium (ERCC) mix (mix 2) was added to a final dilution of 1:25,000,000 within a 10 *µL* volume. The cells went through a standard Smart-Seq2 protocol for mRNA measurement. The stage of ERCC addition makes the absolute values recapitulated directly comparable to the post-lysis mRNA amount.

To preprocess data, all naive and primed cell transcriptomic data was loaded, and forward and reverse reads were summed together. Low read depth cells (*<* 58, 394 reads, the amount of initial transcript features in data frame), were filtered. Read data was collapsed into TPM, normalizing for transcript length. Measured wells with more than 50% of their TPM mapping to ERCCs (control or failed wells) were filtered, resulting in a final count of *n* = 1, 243 single cells.

Expected molecules of each ERCC per well was calculated from known dilution, final well volume, and ERCC mix data, ERCCs with less than 1 expected molecule per cell were ignored. ERCC expected molecules was regressed against ERCC TPM values per cell to find the cell-wise TPM to mRNA molecules conversion factor. That factor, per cell, was then used to scale each TPM value into absolute molecule units, and total endogenous mRNA (post-lysis) was recovered. To reduce transcript features to those consistently expressed, all transcripts that were represented in less than 99% of the cells were removed, resulting in 964 total transcripts being retained. Three cellxgene matrices were then generated for model applications: A absolute molecule count matrix for the linear model, a *log*_2_(*count* + *ϵ*) matrix for the log model, and a proportions matrix for testing. *ϵ* = 3*×*10^*−*4^, corresponding to roughly half the smallest value non-zero value in the molecule counts matrix. Note that molecular counts recapitulation method did not round to integers. Data was split 70-30 for training (*n* = 870) and testing (*n* = 373)

### mRNA model implementation

Both the linear and log-space MCI models were tested across varying feature numbers. The log-space model predictions were implemented in the same way as model 2 of WBC count prediction. Linear-space predictions were able to be done analytically as no *ϵ* offset is needed in linear-space. For all predictions in logarithmic space, an epsilon offset is necessary so that all components of the *c***P** vector are within the domain of the *log*_2_ function. An epsilon of **1** was used to correspond to a single count of the components mRNA.

### mRNA feature selection

Two feature selection methods were tested, in both linear and log-space. Method-one greedily minimizes the predicted RMSE (in respective spaces) of the prediction of mRNA count, as described in the previous method section “Predicting accuracy limits”. This method does not actually need to do the prediction on individual observations to work. In log-space, this method is completely independent of any individual observation’s proportion vector. To implement this in linear-space, a representative proportion vector **P** was used, namely the mean proportion vector of all observations in the training set.

Method 2 is a greedy maximizer which is able to see the resulting predictions of the features it selects. Method 2 maximizes for C-index through 5-fold crossvalidation on each successive feature, and selecting the feature with the highest C-index among average of the 5 cross validation testing sets. Note that all feature selection took place based only on training data, in order to ensure no leakage. The final amount of features to include was chosen through inspection of the four model-method combinations up to 25 features. Note that linear-space predictions were not restricted being positive during feature selection, however the *log*_2_ metrics of RMSE and Pearson’s R were calculated with predicted total mRNA capped at a lower bound of 1. Multiplicative bias is the geometric mean of the predicted values divided by actual, where near 1 values indicate no systemic bias. Gene names in the dataset were originally in Ensemble (ENSG identifier) form and were converted to gene names used biomaRt package[24].

## Data and code availability

Data for mRNA prediction was derived from, and is publicly available at accession E-MTAB-6819 on array express. Flow cytometry and CBC data used to derive absolute counts is available with request to Health and Retirement Study through their distribution system. Code for processing data, analyses, training and testing of models, as well as the model objects with parameters are deposited on Zenodo at 10.5281/zenodo.17513749.

## Author Contributions

J.H. ideated the MCI method, designed and conducted the analyses, derived mathematical formulations, and wrote the manuscript. D.C. reviewed and revised the manuscript with attention to the mathematical accuracy. B.C. provided guidance on analyses plan and method. All authors provided feedback throughout on the manuscript and approve of the final version.

