## Supplementary Information for "Inferring absolute counts from proportions by constraining multivariate normal distributions"

### Supplementary Information for
#### Inferring absolute counts from proportions by constraining multivariate normal distributions

Jeffrey Hage[1], Devin Koestler[2], and Brock Christensen[1,3*]

[1]Department of Epidemiology, Geisel School of Medicine, Dartmouth College, Lebanon, NH
[2]Department of Biostatistics & Data Science, University of Kansas Medical Center, Kansas City, Kansas
[3]Department of Molecular & Systems Biology, Geisel School of Medicine, Dartmouth College, Lebanon, NH
[*]Corresponding author

November 4, 2025

| **Variable** | **Normal** | **$\mathbf{Log_2}$-Normal** | $\boldsymbol{\Delta}$ | **Better fit** |
|---|---|---|---|---|
| WBC | $-1.69 \times 10^4$ | $-1.34 \times 10^4$ | $3.54 \times 10^3$ | log-normal |
| Neu | $-1.20 \times 10^4$ | $-1.15 \times 10^4$ | $5.50 \times 10^2$ | log-normal |
| Bas | $1.16 \times 10^4$ | $1.26 \times 10^4$ | $1.07 \times 10^3$ | log-normal |
| Eos | $2.49 \times 10^3$ | $4.46 \times 10^3$ | $1.97 \times 10^3$ | log-normal |
| Mon | $1.19 \times 10^3$ | $2.00 \times 10^3$ | $8.15 \times 10^2$ | log-normal |
| Bmem | $-1.49 \times 10^4$ | $1.62 \times 10^4$ | $3.12 \times 10^4$ | log-normal |
| Bnv | $3.34 \times 10^3$ | $9.23 \times 10^3$ | $5.90 \times 10^3$ | log-normal |
| CD4mem | $7.97 \times 10^2$ | $1.68 \times 10^3$ | $8.88 \times 10^2$ | log-normal |
| CD4nv | $-1.21 \times 10^3$ | $-5.50 \times 10^1$ | $1.16 \times 10^3$ | log-normal |
| CD8mem | $1.83 \times 10^3$ | $4.56 \times 10^3$ | $2.73 \times 10^3$ | log-normal |
| CD8nv | $8.48 \times 10^3$ | $1.11 \times 10^4$ | $2.58 \times 10^3$ | log-normal |
| NK | $2.45 \times 10^3$ | $4.27 \times 10^3$ | $1.82 \times 10^3$ | log-normal |

**Supplementary Table S1:** Log-likelihood comparison of normal vs. $log_2$-normal (with $\epsilon = 0.003$) curves for WBC and each component cell type with HRS data. Higher log-likelihood indicates a better fit. $\Delta$ column indicates $log_2$-normal minus normal.

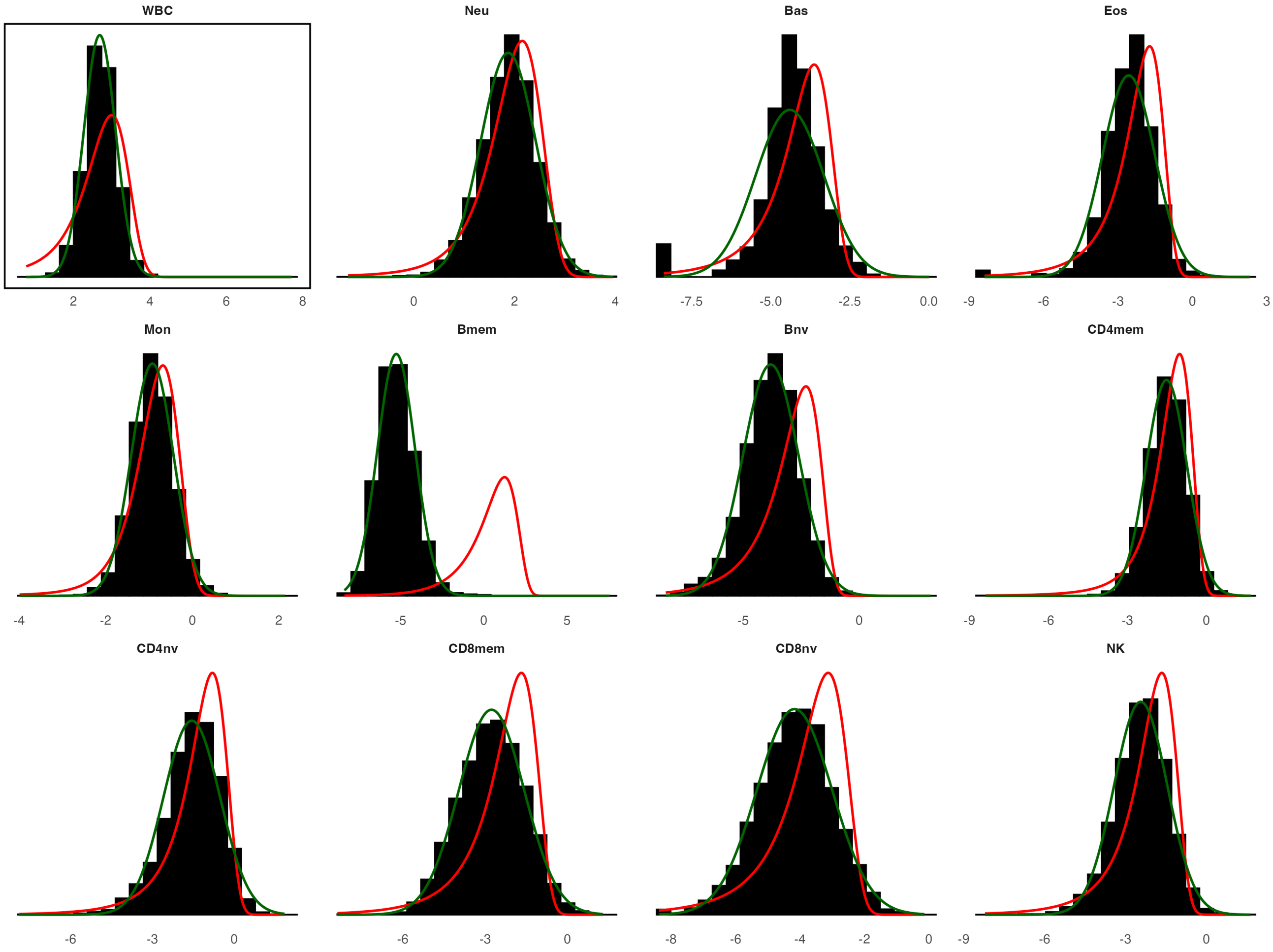


**Supplementary Figure S1:** Distributions of WBC and 11 cell types on $log_2(count + \epsilon)$ with normal (red) and log-normal (green) overlays. X-axis scale is in $log_2$ space.

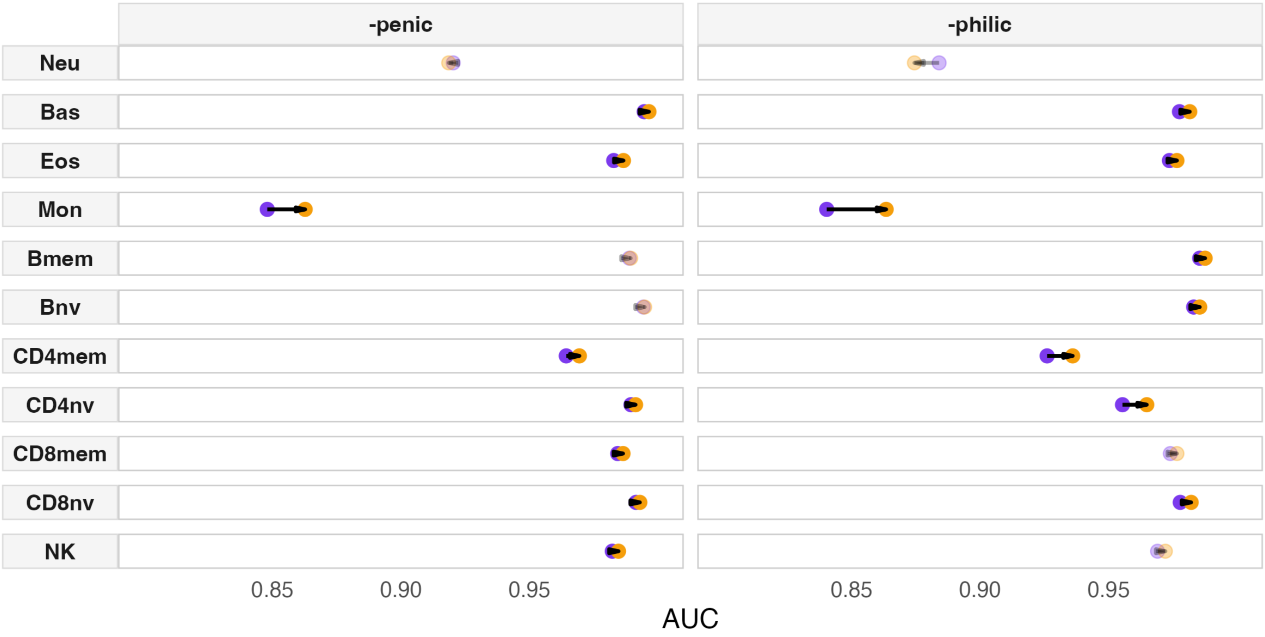


**Supplementary Figure S2:** Classification AUC into cell-type-penic and cell-type-philic within the $n = 6,577$ observations of the health and retirement study. Observations were labeled cell-type-penic or cell-type-philic if they were within the top or bottom 5% of true absolute counts of that cell-type. Purple dots represent the AUC from classification based on proportions, and orange represent from estimated absolute counts. Opaque entries pass FDR$< 0.05$, applied within -penic and -philic groups. Estimate absolute counts significantly (FDR $< 0.05$) outperform true proportions for 16 out of 22 comparisons (adjusted within -penic and -philic seperately), and true proportions did not significantly outperform estimated counts for any cell type.

| **Condition** | **Cell** | **AUC (Abs.)** | **AUC (Est. Abs.)** | **AUC (Prp.)** | **FDR $p$ (ABS vs PRP)** |
|---|---|---|---|---|---|
| **Arthritis** | CD8nv | 0.567 | 0.573 | 0.570 | $7.62 \times 10^{-2}$ |
| | Bmem | 0.545 | 0.548 | 0.549 | $8.03 \times 10^{-1}$ |
| **Diabetes** | CD4mem | 0.579 | 0.546 | 0.535 | $2.20 \times 10^{-6}$ |
| | Neu | 0.580 | 0.539 | 0.528 | $4.49 \times 10^{-3}$ |
| **Alcohol use** | CD8mem | 0.546 | 0.542 | 0.538 | $2.98 \times 10^{-3}$ |
| | CD4mem | 0.534 | 0.526 | 0.521 | $2.79 \times 10^{-2}$ |
| **Lung disease** | Neu | 0.588 | 0.548 | 0.555 | $1.46 \times 10^{-1}$ |
| | Mon | 0.562 | 0.515 | 0.502 | $1.06 \times 10^{-2}$ |

**Supplementary Table S2:** Classification AUC into 4 conditions, by true absolute counts (Abs.), estimated absolute counts (Est. Abs.), and true proportions (Prp.). Estimated absolute counts had significantly (FDR $p < 0.05$) higher AUC than true proportions in 6/8 comparisons, and never the reverse. Cell types comparisons were selected by the two highest AUC of true absolute counts per condition.

| Order Sel. | linear C-index | linear RMSE | log C-index | log RMSE |
|---|---|---|---|---|
| 1 | NDUFB4 | SRP9 | NDUFB4 | **COX7C** |
| 2 | HNRNPH1 | RPL31P2 | NDUFC1 | **SUMO2** |
| 3 | NDUFC1 | KPNA2 | SLC16A1 | **IPO5** |
| 4 | SLIRP | FAM162A | SKP1 | **HSPA4** |
| 5 | BNIP3 | MT-ND3 | RPL15 | **SNRPE** |
| 6 | UBE2D3 | MT-ND6 | ATP5F1B | **HSBP1** |
| 7 | RPA3 | NDUFB4 | RPS27AP5 | **RPL31P2** |
| 8 | SRP9 | HSP90AB3P | CSE1L | **NRDC** |
| 9 | RPL21P119 | PSMD6 | SNRPD1 | **LRPPRC** |
| 10 | PGK1 | HNRNPH1 | LRPPRC | **RPL15** |
| 11 | ZCRB1 | NAP1L1 | SRSF6 | **HSP90AB1** |
| 12 | RHEB | SLC2A1 | GNG5 | **SRP9** |
| 13 | ATP5MG | MT-CYB | TRA2A | **TMSB4X** |
| 14 | FKBP3 | CAP1 | RAD21 | **KDM5B** |
| 15 | TMSB4X | TRMT112 | BTF3L4 | **EIF3M** |
| 16 | PFN2 | HNRNPA1 | SF3B1 | RPL21P119 |
| 17 | LSM3 | NPM1P24 | HSPA8 | MDK |
| 18 | PPP1CB | PCNP | MPLKIP | KPNA2 |
| 19 | HMGN1 | SF3B2 | ATP6V1G1 | DYNLL1 |
| 20 | HMGN3 | IARS1 | RPS2P5 | PTGES3P1 |
| 21 | GAPDHP65 | NDUFC2 | PSMA1 | LARP7 |
| 22 | CTNNB1 | RPL27A | TMEM14C | RPS2P5 |
| 23 | PDCD5 | PRDX6 | DNAJC8 | SNRPB |
| 24 | HMGB1 | BUB3 | TOR1AIP2 | EIF5AL1 |
| 25 | FTLP3 | HSBP1 | NARS1 | YWHAQ |

**Supplementary Table S3:** Selected genes, in order of selection, for each of the four method-model pairs for recapitulating total post-lysis mRNA. Bolded genes correspond to those used in the final mRNA model.

| Cell type | Source | Constituent subtypes | Antibody / gating |
|---|---|---|---|
| B-naive | Flow | N/A | $CD3^-$ $CD19^+$ $IgD^+$ $CD27^-$ |
| B-memory | Flow | Memory B ($IgD^+$)<br>Memory B ($IgD^-$) | $CD3^-$ $CD19^+$ $CD27^+$ $IgD^+$<br>$CD3^-$ $CD19^+$ $CD27^+$ $IgD^-$ |
| CD4-naive | Flow | N/A | $CD3^+$ $CD19^-$ $CD8^-$ $CD4^+$<br>$CD45RA^+$ $CCR7^+$ $CD28^+$ |
| CD4-memory | Flow | CD4 central memory (CM)<br>CD4 effector memory (EM)<br>CD4 TEMRA | Base: $CD3^+$ $CD19^-$ $CD8^-$ $CD4^+$<br>CM: $CD45RA^-$ $CCR7^+$ $CD28^+$<br>EM: $CD45RA^-$ $CCR7^-$ $CD28^-$<br>TEMRA: $CD45RA^+$ $CCR7^-$ $CD28^-$ |
| CD8-naive | Flow | N/A | $CD3^+$ $CD19^-$ $CD8^+$ $CD4^-$<br>$CD45RA^+$ $CCR7^+$ $CD28^+$ |
| CD8-memory | Flow | CD8 central memory (CM)<br>CD8 effector memory (EM)<br>CD8 TEMRA | Base: $CD3^+$ $CD19^-$ $CD8^+$ $CD4^-$<br>CM: $CD45RA^-$ $CCR7^+$ $CD28^+$<br>EM: $CD45RA^-$ $CCR7^-$ $CD28^-$<br>TEMRA: $CD45RA^+$ $CCR7^-$ $CD28^-$ |
| Natural killer | Flow | N/A | $CD3^-$ $CD19^-$ $CD20^-$ $CD14^-$<br>$CD16^+$ $CD56^+$ |
| Monocytes | CBC | N/A | N/A |
| Neutrophils | CBC | N/A | N/A |
| Eosinophils | CBC | N/A | N/A |
| Basophils | CBC | N/A | N/A |

**Supplementary Table S4:** Table describing how each of the 11 cell types used for absolute count prediction were derived from HRS data, and the antibody scheme used in flow cytometry when applicable. B-memory, CD4-memory, and CD8-memory were derived via collapsing multiple absolute counts. Monocyte and dendritic flow cytometry data was not used.
